# Human gut metagenomes and assembled microbial genomes from an urban cohort from Colombia, South America

**DOI:** 10.1101/2025.09.25.678255

**Authors:** Jacobo de la Cuesta-Zuluaga, Nicholas D. Youngblut, Ruth E. Ley, Juan S. Escobar

## Abstract

This study presents shotgun gut metagenomes of 430 men and women (18-62 years old) from diverse urban areas of Colombia. Metagenome assembly resulted in 2 266 medium- and high-quality metagenome-assembled genomes (MAGs). The metagenomes and MAGs complement previously published 16S rRNA gene and host data from the same cohort.

## Announcement

The study of the human gut microbiome has transformed the interpretation of multiple physiological processes, from health and disease to metabolism and nutrient absorption (1). However, large-scale studies from low- and middle-income countries aiming to describe gut microbial diversity and its association with human health are sparse (2). The lack of studies in such populations makes it difficult to determine the generality of many of the previously reported links between the microbiome and the host.

Here, we report a set of shotgun metagenomes and metagenome-assembled genomes (MAGs) from the gut of 430 community-dwelling human adults from five cities in Colombia, South America. These subjects were enrolled as part of a research project aimed at characterizing the gut microbiota of a population undergoing westernization and to determine variation associated with obesity and cardiometabolic disease (3). The metagenome sequencing data were used to validate previously reported features of gut microbiome function and diversity independently correlated with obesity or the cardiometabolic health of the hosts (4).

Total DNA was extracted from fecal samples utilizing the QIAamp DNA Stool Mini Kit (Qiagen, Hilden, Germany). For the newly generated metagenomes, we prepared shotgun metagenome libraries with a modified Nextera protocol (5). We used 1 ng of total stool DNA for Nextera Tn5 tagmentation. After purification with Agencourt AMPure XP beads (Beckman Coulter, Brea, CA, USA), samples were normalized and pooled. We performed size selection of the pooled samples using BluePippin (Sage Sciences, Beverly, MA, USA) to restrict fragment sizes to 400 to 700 bp. Barcoded pools were sequenced using the Illumina HiSeq 3000 platform with 2x150 bp paired-end sequencing. We included blank DNA extraction samples as negative controls and DNA from synthetic mock communities, as well as fecal samples from a previously sequenced donor, as positive controls. Library preparation and sequencing were performed at the Max Planck Institute for Biology Tübingen, Germany, and the use of the metagenomes was reported by de la Cuesta-Zuluaga et al. (4).

Raw sequencing reads were validated using fqtools v.2.0 (6), and the “clumpify” command of bbtools v37.78 (https://jgi.doe.gov/data-and-tools/bbtools/) was used to deduplicate them. We removed adapters and performed read quality control with skewer v0.2.2 (7) and the “bbduk” command of bbtools. We removed human reads using the “bbmap” command of bbtools against the hg19 genome assembly. We generated quality reports using fastqc v0.11.7 (https://github.com/s-andrews/FastQC) and multiQC v1.5a (8). Metagenome coverage was calculated using Nonpareil v.3.3.4 (9). We retained 408 samples with a sequencing depth of >5.0x10^5^ reads (mean ± SD 6 719 985 reads/sample ± 8 960 996) or a nonpareil coverage > 60 % (82.36 % ± 8.38) (Figure 1A).

**Figure 1.**
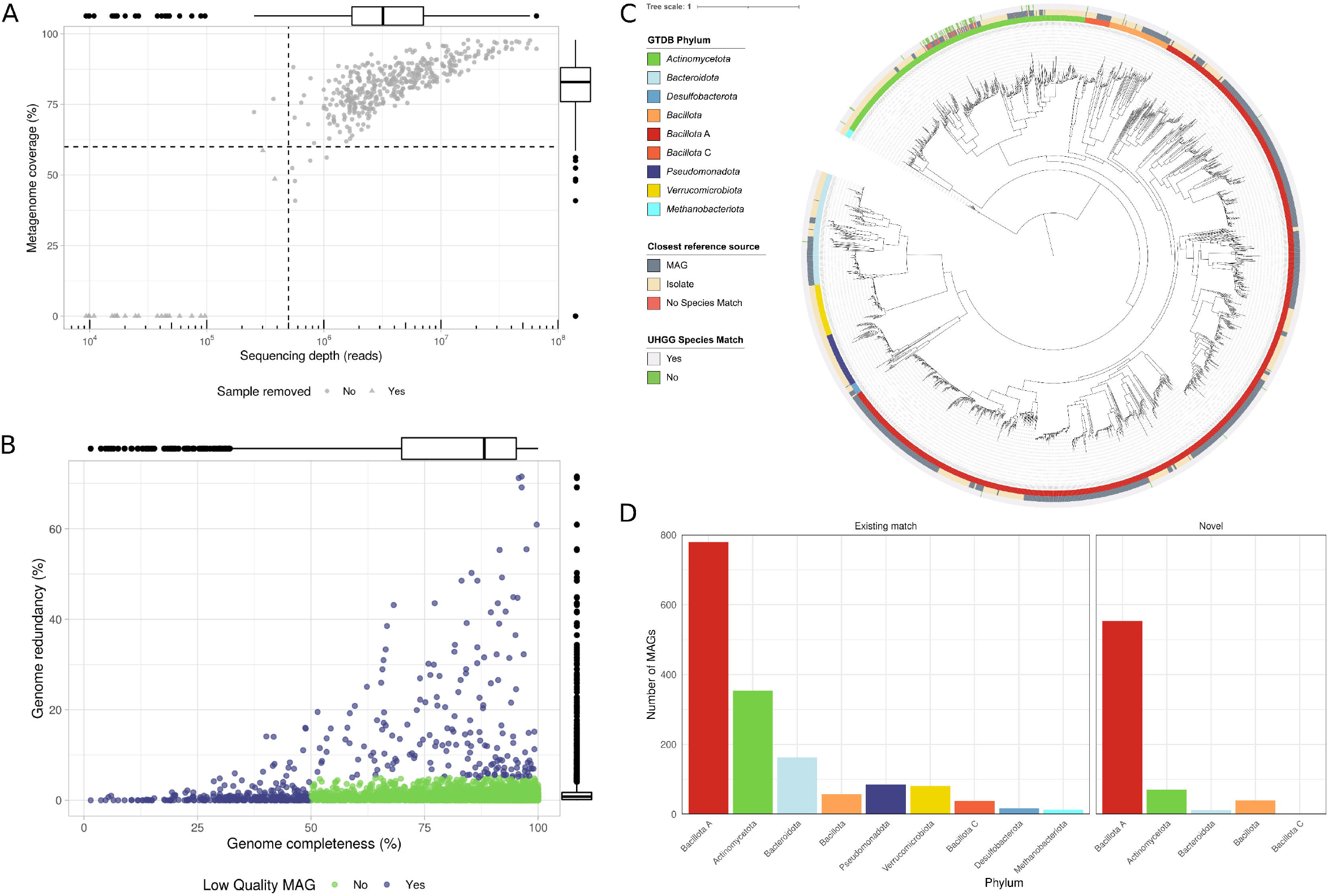
Summary of shotgun metagenome sequencing and metagenome assembly from 430 human gut samples. A) Sequencing depth and metagenome coverage of metagenome samples; dashed lines represent inclusion thresholds for each parameter. B) Completeness and redundancy of each of the obtained MAGs. Green points correspond to the 2 266 MAGs above quality cut-off values. C) Phylogenetic tree built from multilocus sequence alignment. From innermost to outermost rings, the data mapped onto the phylogeny are: GTDB r214 phylum-level taxonomic classification; the source of the genome of the closest match to the GTDB representative set (environmental sample: MAG, a cultured isolate: Isolate; the Colombian MAG had no species-level match); presence of a genome with ANI < 95% at the UHGG catalog v.1.0. The phylogeny was inferred from multiple conserved loci using PhyloPhlAn. The phylogeny is rooted on the last common ancestor of *Archaea* and *Bacteria*. The scale bar represents the number of amino acid substitutions per site. D) Barplot showing the number of MAGs classified on each bacterial and archaeal phylum according to whether they belonged to genera with a valid name (Existing match), or whether they did not have a species-level classification or belonged to a genus with no binomially named representative genome in the GTDB (Novel).

We performed metagenome assembly following the workflow developed by Youngblut et al. (10): each sample was assembled separately, prior to subsampling to a maximum of 20 million reads per sample using seqtk v.1.3. A reference-based metagenome assembly was performed using metacompass v.1.2 (11) based on each sample’s taxonomic profile. We profiled each sample using centrifuge v.1.0.3 (12) to select reference genomes, which we then downloaded using ncbi-genome-download v.0.2.1. Reads that did not map to any reference genome were used for de-novo assembly with metaSPAdes v.3.12.0 (13). Contigs with a minimum length of 2 000 bp from the reference-based and de novo assemblies were combined and de-replicated using bbtools. To bin contigs, we used MaxBin2 v.2.2.4 (14) and MetaBAT2 v.2.12.1 (15), each executed with 2 parameter settings for a total of 4 bin collections per sample. Per-sample binning of contigs was performed, and we utilized reads from all metagenome samples to calculate differential coverage with Bowtie2 v.2.3.5 (16). The best non-redundant set of contig bins (i.e., MAGs) was selected with DAS-Tool v.1.1 (17) based on estimates of completeness and contamination from CheckM (18).

We calculated completeness and contamination of each MAG using CheckM v.1.0.13 (18). MAGs with completeness of < 50 % or contamination of ≥ 5 % were discarded. Taxonomic classification of the MAGs was obtained using GTDB-Tk v.0.3.3 (19) against release 214 of the Genome Taxonomy Database (GTDB). We used dRep v. 2.5.4 (https://github.com/MrOlm/drep) to collapse clonal genomes at an average nucleotide identity (ANI) of 99.9 %. We retrieved 2 797 MAGs; after removal of low-quality genomes and dereplication, we retained 2 266 MAGs, with an average completeness and contamination of 85.57 % ± 13.11 and 1.07 % ± 1.04, respectively (Figure 1B).

Using PhyloPhlAn v.0.41 (20), we constructed a maximum-likelihood phylogeny of de-replicated MAGs (Figure 1C). We assessed the taxonomic novelty of the non-redundant MAGs against the Unified Human Gut Genomes catalog v.1.0 (UHGG) (21) using FastANI v.1.31 (22). We considered a MAG to be novel if it did not have a species-level match in the UHGG (< 95 % ANI) (Figure 1D).

## Data availability

The raw metagenomic sequence data generated during this study and the corresponding non-redundant metagenome-assembled genomes have been deposited in the European Nucleotide Archive with accession ID PRJEB58436. Host anthropometric, biochemical, and dietary data have been made available as part of previously published works and can be found in the following repositories: https://github.com/jsescobar/westernization, https://github.com/jsescobar/bsp, and https://github.com/jsescobar/diet_microbiota_MiSalud1.0. The code used for processing the data can be found at https://github.com/jacodela/Colombian_MAGs.

## Competing Interests

While engaged in the research project, J.S.E. was employed by Vidarium, a research center belonging to a food company (Grupo Empresarial Nutresa). J.d.l.C-Z., N.D.Y., and R.E.L. declare no competing interests.

## Acknowledgements

This work was supported by the Max Planck Society (J.d.l.C-Z., N.D.Y., R.E.L.) and Vidarium–Nutrition, Health and Wellness Research Center (J.S.E.). We thank the participants who took part in the study, and the Vidarium, EPS SURA, and Dinámica IPS staff who helped with recruitment and field work. Some authors of this work collaborate through the Microbiome & Health Network. We are grateful to Liam Fitzstevens, Taichi Suzuki, and Laura Salazar-Jaramillo for the fruitful discussions and comments.

